# Long-reads assembly of the *Brassica napus* reference genome, Darmor-bzh

**DOI:** 10.1101/2020.07.22.215749

**Authors:** Mathieu Rousseau-Gueutin, Caroline Belser, Corinne Da Silva, Gautier Richard, Benjamin Istace, Corinne Cruaud, Cyril Falentin, Franz Boideau, Julien Boutte, Regine Delourme, Gwenaëlle Deniot, Stefan Engelen, Julie Ferreira de Carvalho, Arnaud Lemainque, Loeiz Maillet, Jérôme Morice, Patrick Wincker, France Denoeud, Anne-Marie Chèvre, Jean-Marc Aury

**Author notes:** Equal contribution (Mathieu Rousseau-Gueutin, Caroline Belser, Corinne Da Silva, Gautier Richard, Benjamin Istace, Corinne Cruaud, Cyril Falentin).

## Abstract

**Background:** The combination of long-reads and long-range information to produce genome assemblies is now accepted as a common standard. This strategy not only allow to access the gene catalogue of a given species but also reveals the architecture and organisation of chromosomes, including complex regions like telomeres and centromeres. The *Brassica* genus is not exempt and many assemblies based on long reads are now available. The reference genome for *Brassica napus*, Darmor-bzh, which was published in 2014, has been produced using short-reads and its contiguity was extremely low if compared to current assemblies of the *Brassica* genus.

**Findings:** Here, we report the new long-reads assembly of Darmor-bzh genome (*Brassica napus*) generated by combining long-reads sequencing data, optical and genetic maps. Using the PromethION device and six flowcells, we generated about 16M long-reads representing 93X coverage and more importantly 6X with reads longer than 100Kb. This ultralong-reads dataset allows us to generate one of the most contiguous and complete assembly of a *Brassica* genome to date (contigs N50 > 10Mb). In addition, we exploited all the advantages of the nanopore technology to detect modified bases and sequence transcriptomic data using direct RNA to annotate the genome and focus on resistance genes.

**Conclusion:** Using these cutting edge technologies, and in particular by relying on all the advantages of the nanopore technology, we provide the most contiguous *Brassica napus* assembly, a resource that will be valuable for the *Brassica* community for crop improvement and will facilitate the rapid selection of agronomically important traits.

## Background

Recently, Pacific Biosciences (PACBIO) and Oxford Nanopore (ONT) sequencing technologies were commercialized with the promise to sequence long DNA fragments (kilobases to megabases order). The introduction of these long-reads technologies increased significantly the quality in terms of contiguity and completeness of genome assemblies [1–3]. Although it is possible to assemble chromosomes into a single contig in the case of simple genomes [4], the complexity of plant genomes is such that these methods are still insufficient to obtain the chromosome architecture. The resulting contigs need to be ordered and oriented with long-range information based on chromatin interactions or optical maps. This combined strategy is now a common standard.

*Brassica* crops include important vegetables for human nutrition and for production of vegetable oil. It is composed of several well studied species that belong to the U’s triangle: three diploid species *B. rapa* (AA), *B. nigra* (BB) and *B. oleracea* (CC) and three allotetraploid hybrid species *B. juncea* (AABB), *B. napus* (AACC) and *B. carinata* (BBCC). As *Brassica* species underwent several polyploidy events, the genomes of the diploid and allopolyploid species are important models to understand the immediate and long term impact of polyploidy on the structural and the functional evolutionary dynamic of duplicated genes and genomes. These events had a significant role in their diversification, indeed the variability between two morphotypes of the same *Brassica* species can be high, underlining the importance of having a collection of high quality assemblies for the *Brassica* genus and more generally for all species.

To date, several genomes of the *Brassica* genus have been sequenced using long-reads and brought to the chromosome scale. The two first ones, *B. rapa* Z1 and *B. oleracea* HDEM, were published in 2018 and generated using ONT reads and optical maps [2,4]. At the beginning of 2020, two genomes of *B. rapa* (Chiifu) and *B. oleracea* (D134) have been released and sequenced using PACBIO long reads [5][6]. A first long-reads assembly based on ONT of the other diploid species, *B. nigra* NI100 is available since the beginning of 2020 [7]. The availability of the diploid genomes first may be explained by their smaller genome size (<600Mb) compared to their tetraploid derivatives. The first genome of a tetraploid plant has been published in 2014 [8] and was sequenced using a short-reads strategy (referred here as *B. napus* Darmor-bzh v5), and although a great resource for the scientific community, this genome remains very fragmented and incomplete. In 2017, the pseudomolecules of Darmor-bzh were improved [9] using GBS data (referred here as Darmor-bzh v8), and a new gene prediction was made. Unfortunately, this annotation was not used by the Brassica community as it was considered incomplete [10]. As an illustration, only 89.6% of the core brassicale genes were found in the annotation compared to the 97.7% of the previous version (see Table 1). In the first months of 2020, nine *B. napus* genomes based on PACBIO sequencing were published [11,12], and three of them were organized using long-range information. These genomes have a better continuity than the current reference (Darmor-bzh), but interestingly their N50 at the contig level is lower than the *Brassica* genomes that have been generated using ONT sequencing (Table 2). Here, we report the genome sequence of the *B. napus* reference Darmor-bzh (referred here as Darmor-bzh v10), produced using ONT and Illumina reads supplemented by optical and genetic maps. In addition to chromosomes architecture, genes were predicted by sequencing native RNA molecules using the ONT PromethION with a focus on resistance genes. The contiguity and gene completion of this new genome assembly is among the highest in the *Brassica* genus (Table 2).

**Table 1.** Statistics of the Darmor-bzh assemblies.

|  | version | technology | contigs<br>N50 (L50) | #<br>contigs | Cumulative<br>size | Number<br>of genes | Number<br>of<br>anchored<br>bases | Number<br>of<br>anchored<br>[ACGT]<br>bases | BUSCO<br>Complete<br>n=4596 |
| --- | --- | --- | --- | --- | --- | --- | --- | --- | --- |
| *B.*<br>*napus* | 10 | ONT | 11,486,274 (24) | 505 | 924 Mb | 108,190 | 867 Mb<br>(93.84%) | 849 Mb | 98.6% |
| *B.*<br>*napus* | 8 | 454 | 37,644<br>(5,517) | 44,818 | 850 Mb | 80,382 | 799 Mb<br>(93.96%) | 690 Mb | 89.6% |
| *B.*<br>*napus* | 5 | 454 | 37,644<br>(5,517) | 44,837 | 850 Mb | 101,040 | 645 Mb<br>(75.90%) | 553 Mb | 97.7% |

**Table 2.**
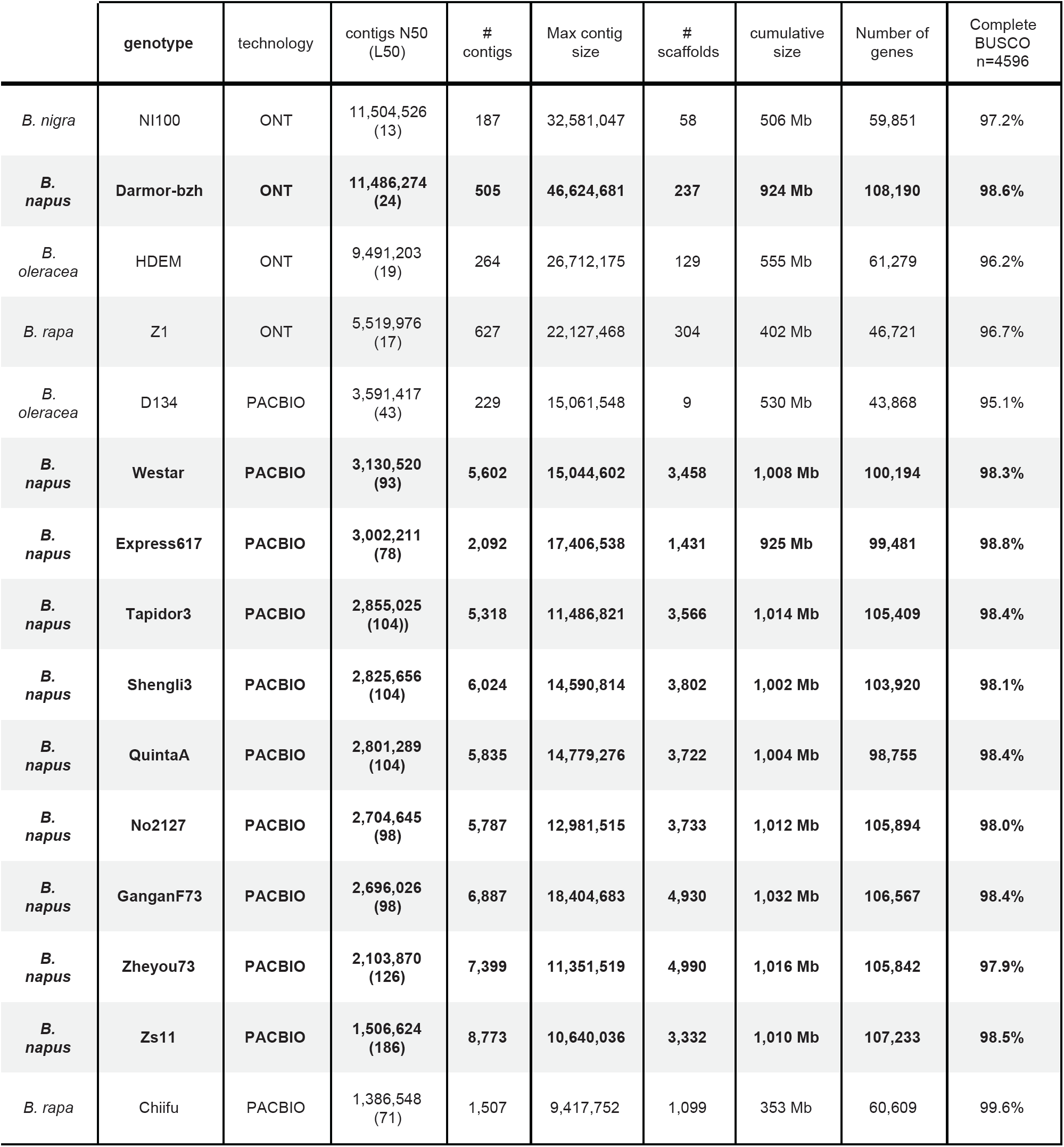
Statistics of the *Brassica* long-reads assemblies (ordered by contigs N50 value, *B. napus* are in bold)

|  | genotype | technology | contigs N50 (L50) | # contigs | Max contig size | # scaffolds | cumulative size | Number of genes | Complete BUSCO n=4596 |
| --- | --- | --- | --- | --- | --- | --- | --- | --- | --- |
| *B. nigra* | NI100 | ONT | 11,504,526 (13) | 187 | 32,581,047 | 58 | 506 Mb | 59,851 | 97.2% |
| ***B. napus*** | **Darmor-bzh** | **ONT** | **11,486,274 (24)** | **505** | **46,624,681** | **237** | **924 Mb** | **108,190** | **98.6%** |
| *B. oleracea* | HDEM | ONT | 9,491,203 (19) | 264 | 26,712,175 | 129 | 555 Mb | 61,279 | 96.2% |
| *B. rapa* | Z1 | ONT | 5,519,976 (17) | 627 | 22,127,468 | 304 | 402 Mb | 46,721 | 96.7% |
| *B. oleracea* | D134 | PACBIO | 3,591,417 (43) | 229 | 15,061,548 | 9 | 530 Mb | 43,868 | 95.1% |
| ***B. napus*** | **Westar** | **PACBIO** | **3,130,520 (93)** | **5,602** | **15,044,602** | **3,458** | **1,008 Mb** | **100,194** | **98.3%** |
| ***B. napus*** | **Express617** | **PACBIO** | **3,002,211 (78)** | **2,092** | **17,406,538** | **1,431** | **925 Mb** | **99,481** | **98.8%** |
| ***B. napus*** | **Tapidor3** | **PACBIO** | **2,855,025 (104)** | **5,318** | **11,486,821** | **3,566** | **1,014 Mb** | **105,409** | **98.4%** |
| ***B. napus*** | **Shengli3** | **PACBIO** | **2,825,656 (104)** | **6,024** | **14,590,814** | **3,802** | **1,002 Mb** | **103,920** | **98.1%** |
| ***B. napus*** | **QuintaA** | **PACBIO** | **2,801,289 (104)** | **5,835** | **14,779,276** | **3,722** | **1,004 Mb** | **98,755** | **98.4%** |
| ***B. napus*** | **No2127** | **PACBIO** | **2,704,645 (98)** | **5,787** | **12,981,515** | **3,733** | **1,012 Mb** | **105,894** | **98.0%** |
| ***B. napus*** | **GanganF73** | **PACBIO** | **2,696,026 (98)** | **6,887** | **18,404,683** | **4,930** | **1,032 Mb** | **106,567** | **98.4%** |
| ***B. napus*** | **Zheyu73** | **PACBIO** | **2,103,870 (126)** | **7,399** | **11,351,519** | **4,990** | **1,016 Mb** | **105,842** | **97.9%** |
| ***B. napus*** | **Zs11** | **PACBIO** | **1,506,624 (186)** | **8,773** | **10,640,036** | **3,332** | **1,010 Mb** | **107,233** | **98.5%** |
| *B. rapa* | Chiifu | PACBIO | 1,386,548 (71) | 1,507 | 9,417,752 | 1,099 | 353 Mb | 60,609 | 99.6% |

## DNA extraction and sequencing

### Plant material

*B. napus* cv. Darmor-bzh is a French winter variety, freely available in Plant Genetic Resources BrACySol (Rennes, France). Darmor-bzh seeds were sown in Fertiss blocks (Fertil, France). Plantlets were grown under 16h light/8h night in a greenhouse at 20°C for 10 to 12 days of culture. Prior to harvest, plants were either dark treated during 5 days for DNA extraction or not treated after the 10 to 12 days of culture for RNA extraction.

### DNA and RNA extractions

*B. napus* Darmor-bzh DNA extractions for nanopore sequencing were prepared from 1cm2 of first young leaves from which the midribs were removed. Samples were placed on an aluminium foil on ice. 2.5g of each genotype were N2 ground with mortar and pestle during 1 minute. Ground materials were homogenized with 10ml pre-heated CF lysis buffer (MACHEREY-NAGEL GmbH & Co. KG) supplemented with 4mg proteinase K in 50ml tubes containing phase-lock gel and incubated for 45min at 56°C. 10ml of saturated phenol (25:24:1) was added to samples and tubes were placed on a rotator at 40 rpm for 10min to get a fine emulsion. Samples were centrifuged for 24min at 4500g (Acc3/Dec3) and aqueous phases were poured in new 50ml tube containing phase-lock gel. 10ml of chloroform-octanol (24:1) were added to each sample and tubes were placed on a rotator at 40 rpm for 10min to get a fine emulsion. Samples were centrifuged for 24min at 4500g (Acc3/Dec3) and aqueous phases were poured in new 50ml tube. DNA precipitation was done by adding 4ml of NaCl 5M and 30ml of cold 100% isopropanol. After 3 hours at 4°C, DNA was hooked out in one-piece with a hook by melting a glass capillary in a blue flame. DNA pellets were submerged in a 50 ml tube containing 70% ethanol and transferred in a new tube to evaporate remaining isopropanol at 37°C (oven). Dried DNA was resuspended with 3ml TE 10/1 buffer. Quality of extraction was evaluated using a 2200 TapeStation System (Agilent Technologies) with a Genomic DNA kit.

Total RNA was extracted from 2.5g *B. napus* leaf and root tissues using LiCl modified protocol [13]. RNA was controlled using a Bioanalyzer Chip (Agilent RNA 6000 Pico Kit) and quantified using Qubit (RNA HS Assay Kit).

### Illumina Sequencing

DNA (1.5μg) was sonicated to a 100-to 1500-bp size range using a Covaris E220 sonicator (Covaris, Woburn, MA, USA). Fragments (1µg) were end-repaired, 3′-adenylated and Illumina adapters were then added using the Kapa Hyper Prep Kit (KapaBiosystems, Wilmington, MA, USA). Ligation products were purified with AMPure XP beads (Beckman Coulter Genomics, Danvers, MA, USA). Libraries were then quantified by qPCR using the KAPA Library Quantification Kit for Illumina Libraries (KapaBiosystems), and library profiles were assessed using a DNA High Sensitivity LabChip kit on an Agilent Bioanalyzer (Agilent Technologies, Santa Clara, CA, USA). Libraries were sequenced on an Illumina HiSeq2500 instrument (Illumina, San Diego, CA, USA) using 250 base-length read chemistry in a paired-end mode.

After the Illumina sequencing, an in-house quality control process was applied to the reads that passed the Illumina quality filters as described in [14]. These trimming and removal steps were achieved using in-house-designed software based on the FastX package [13–15]. This processing resulted in high-quality data and improvement of the subsequent analyses (Table S1).

### Nanopore Sequencing (PromethION)

The libraries were prepared according to the protocol Genomic DNA by ligation (SQK-LSK108 and SQK-LSK109 Kit) – PromethION’ provided by Oxford Nanopore. Genomic DNA was first repaired and End-prepped with the NEBNext FFPE Repair Mix (New England Biolabs, Ipswich, MA, USA) and the NEBNext® Ultra™ II End Repair/dA-Tailing Module (NEB). DNA was then purified with AMPure XP beads (Beckman Coulter, Brea, CA, USA) and sequencing adapters provided by Oxford Nanopore Technologies were ligated using Concentrated T4 DNA Ligase 2M U/ml (NEB). After purification with AMPure XP beads (Beckman Coulter) using Dilution Buffer (ONT) and Wash Buffer (ONT), the library was mixed with the Sequencing Buffer (ONT) and the Library Loading Bead (ONT), and loaded on the PromethION Flow Cells (R9.4.1).

No data cleaning was performed, the nanopore long reads were used raw for all the assemblies (Table S1). A taxonomical assignation was performed using Centrifuge [16] for each dataset to detect potential contaminations.

### Direct RNA sequencing (PromethION)

rRNA was depleted using the Ribo-Zero rRNA Removal Kit (Plant Leaf). RNA libraries were performed from a mix of 960 ng mRNA according to the ONT protocol “Direct RNA sequencing” (SQK-RNA002). We performed the optional reverse transcription step to improve throughput, but cDNA strand was not sequenced. The Nanopore library was sequenced using the PromethION on a single flowcell (Table S9).

### Optical Maps

*B.napus* Darmor-bzh DNA extractions for nanopore sequencing were prepared from 1cm2 of first young leaves from which the midribs were removed. Samples were placed on an aluminium foil on ice. 5g of each genotype were N2 ground with mortar and pestle during 2 minutes. Ground materials were homogenized in 50ml NIBTM (10mM Tris-HCl pH 8.0, 10mM EDTA pH 8.0, 80mM KCl, 0.5M Sucrose, 1mM Spermine tetrahydrochloride, 1mM spermidine trihydrochloride, 2% (w/v) PVP40, pH was adjusted to 9.4 and solution was filtered through 0.22µm (NIB) and supplemented with 0.5% TritonX-100 (NIBT) and 7.5% 2-Mercaptoethanol (NIBTM). Nuclei suspensions were filtered through Cheese Cloth and Mira Cloth and centrifuged at 1500g during 20min at 4°C. Pellets were suspended in 1ml NIBTM and adjusted to 20ml with NIBTM. Nuclei suspensions were filtered again through Cheese Cloth and Mira Cloth and centrifuged at 57g during 2min at 4°C. Supernatants were kept and centrifuged at 1500g during 20min at 4°C. Pellets were suspended in 1ml NIBT and adjusted to 20ml with NIBT. In order to wash the pellets, the last steps were repeated 3 times with 50ml of NIBT and a final time with 50ml of NIB. Pellets were suspended in residual NIB (approximately 200µl), transferred in a 1.5ml tube and centrifuged at 1500g during 2min at 4°C. Nuclei were suspended in cell suspension buffer from CHEF Genomic DNA Plug Kits (Bio-Rad) and melted agarose 2% from the same kit was added to reach a 0.75% agarose plug concentration. Plug lysis and DNA retrieval were performed as recommended by Bionano Genomics.

The NLRS labeling (BspQI) protocol was performed according to Bionano with 600ng of DNA. The DLS labeling (DLE-1), was performed with 750ng of DNA. Loading of the chip was performed as recommended by Bionano Genomics.

### *B. napus* Darmor-bzh genetic map

An integrated *B. napus* genetic map was obtained using BioMercator V4.2 program [17] from three different genetic maps [18] : a cross between Darmor-bzh and Yudal (DH population with 248 lines), a cross between Darmor and Bristol (F2 population with 291 plants) and a cross between Darmor and Samouraï (DH population with 129 lines). These populations were genotyped using the Illumina 8K, 20K and 60K arrays [18][8][19][20] and genetic maps were constructed using CarthaGene software [21].

## Genome assembly

### Long reads genome assembly

In order to get the best possible genome assembly, we used three assemblers : redbean [22], SMARTdenovo [23] and Flye [24] with all nanopore reads or subsets of reads composed of either the longest reads or those selected by the Filtlong [25] software with default parameters (Table S2), as it has been proven that it could be beneficial for the assembly phase to downsample the reads coverage [3]. We used the following options as input to SMARTdenovo : “-c 1” to generate a consensus sequence, “-J 5000” to remove sequences smaller than 5kb and “-k 17” to use 17-mers as it is advised by developers in the case of large genomes.

Then, we selected the ‘best’ assembly based not only on contiguity metrics such as N50 but also cumulative size. The Flye assembler using the longest reads produced the most contiguous assembly and lead to a contig N50 of 10.0Mb.

As nanopore reads contain systematic error in homopolymeric regions, we polished the consensus of the selected assembly three times with nanopore reads as input to the Racon [26] software and then three additional times using Illumina reads as input to the Pilon tool [27]. Both tools were used with default parameters (Table S6).

### Long range genome assembly

The genome map assemblies of *B. napus* Darmor-bzh have been generated using the Bionano Solve Pipeline version 3.3 and the Bionano Access version 1.3 (Tables S3-S5). The assemblies were performed using the parameters “non haplotype without extend and split” and “add Pre-assembly”. This parameter allows to obtain a rough assembly used as reference for a second assembly. We filtered out molecules smaller than 180Kb and molecules with less than nine labeling sites (Tables S3 and S4). The Bionano scaffolding workflow was launched with the nanopore contigs and the two Bionano maps (DLE and BspQI). As already reported, we found in several cases that the nanopore contigs were overlapping (based on the optical map) and this overlap was not managed by the hybrid scaffolding procedure. We corrected this negative gaps using the BiSCoT software [28] with default parameters (Table S6).

### Validation and anchoring of the *B. napus* Darmor-bzh assembly

A total of 39,495 markers deriving from the Illumina 8K, 20K and 60K arrays were genetically mapped on the integrated map, totalling 2,777.7 cM. The sequence contexts of all SNP markers that were genetically mapped were blasted against our *B. napus* Darmor-bzh assembly in order to validate the quality of our assembly and to help ordering and orientating the scaffolds. Of these 39,495 genetically mapped markers, 36,391 were physically anchored on the final Darmor-bzh assembly (Figure 1 and Table S7). The genetic and physical positions were discordant for only 618 markers (0.02%) due to an inaccurate position on the genetic map (variation of a few centimorgans in almost all cases).

**Figure 1.**
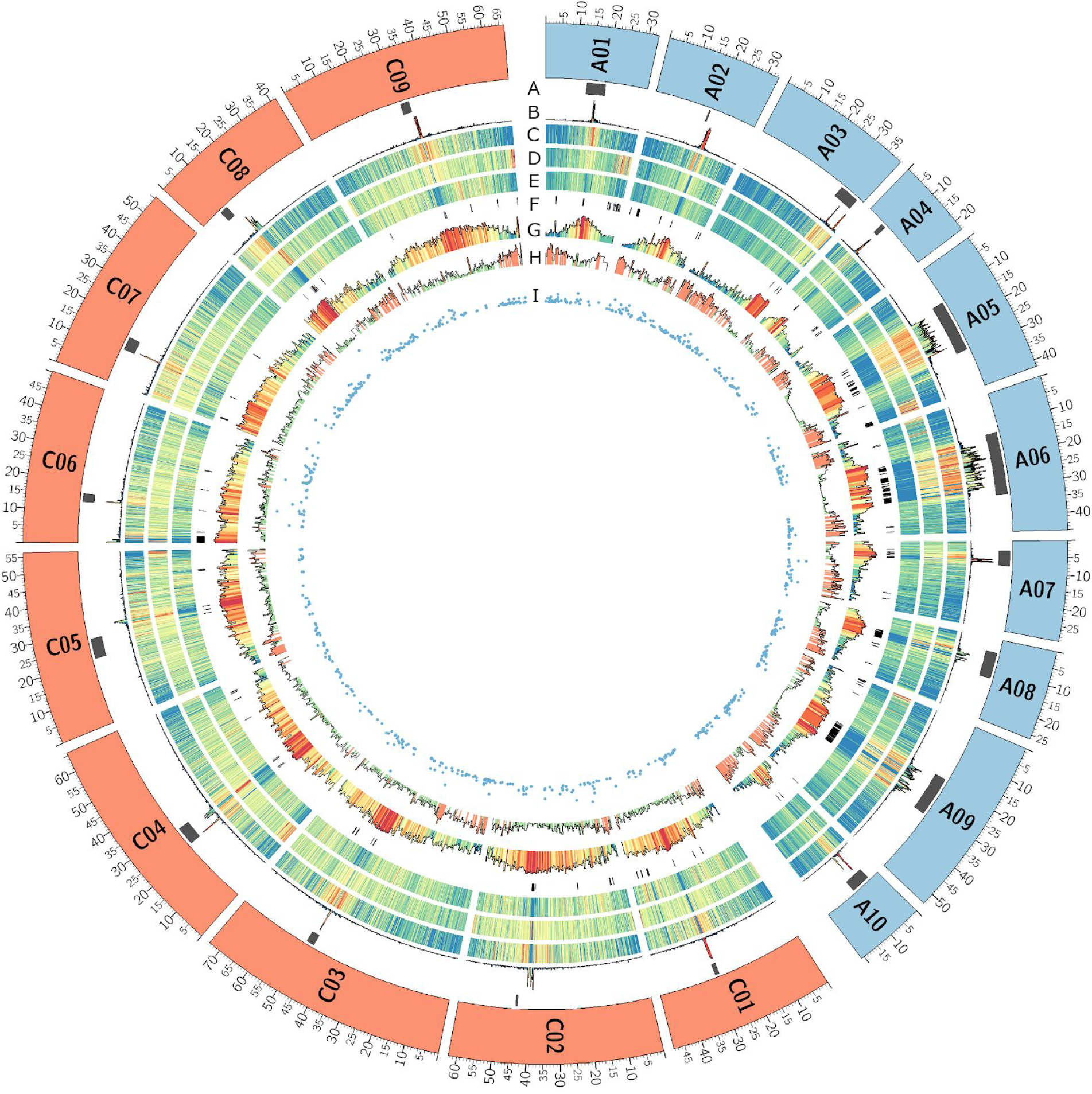
Genome overview of the 19 chromosomes of *B. napus* Darmor-bzh v10 (the ten A chromosomes are in blue and the nine C chromosomes in orange). **A**. Localization of centromere from flanking markers identified by [45], **B**. Density of repeated pericentromeric-specific sequences of *Brassica* allowing more precise localization of centromeres. **C**. Density of Gypsy elements. **D**. Density of Copia elements. **E**. Density of DNA transposon elements. **F**. black boxes represent gaps in the Darmor-bzh assembly. **G**. Density of methylated CpG. **H**. Gene density. **I**. scatter plots of RGAs density. All densities are calculated in 100Kb windows, blue and red colors in density plots are for lower and higher values respectively.

### Identification of large misassembled inverted regions

In order to compare this new *B. napus* Darmor-bzh assembly (v10) to the first published version (v5) as well as to the recent *B. napus* ZS11 genome assembly [11], we performed whole chromosome alignments using nucmer from Mummer [29], followed by automatic plotting using ggplot in R [30]. Nucmer was used with default parameters, followed by delta-filter with the ′-i 98 -l 1000′ parameters. Using the output of Nucmer and delta-filter described above, we subsequently identified Large Misassembled Inverted (LMI) regions between the v5 and v10 genome assemblies. We used show-coords from Mummer with the following parameters ′-H -d -r -T′, followed by a specific method to extract LMIs. Since the show-coords output is consisting in many small alignments from which we need to infer LMIs, we used an iterative process based on merging the coordinates of small consecutive inversions in larger ones, followed by filtering out the small inversions. The parameters were validated by visually and manually checking the resulting LMIs coordinates against the chromosome by chromosome alignment scatter plots (Figure 2). More precisely, we extracted LMIs from the show-coords output, followed by ‘bedtools merge -d 100000’ [31] to merge the coordinates of inversions separated by 100Kb or less, then we selected inversions of size superior to 100Kb, subsequently merged the inversions separated by 300Kb with bedtools, selected inversions larger than 700Kb, and finally merged inversions separated by 1Mb with bedtools.

**Figure 2.**
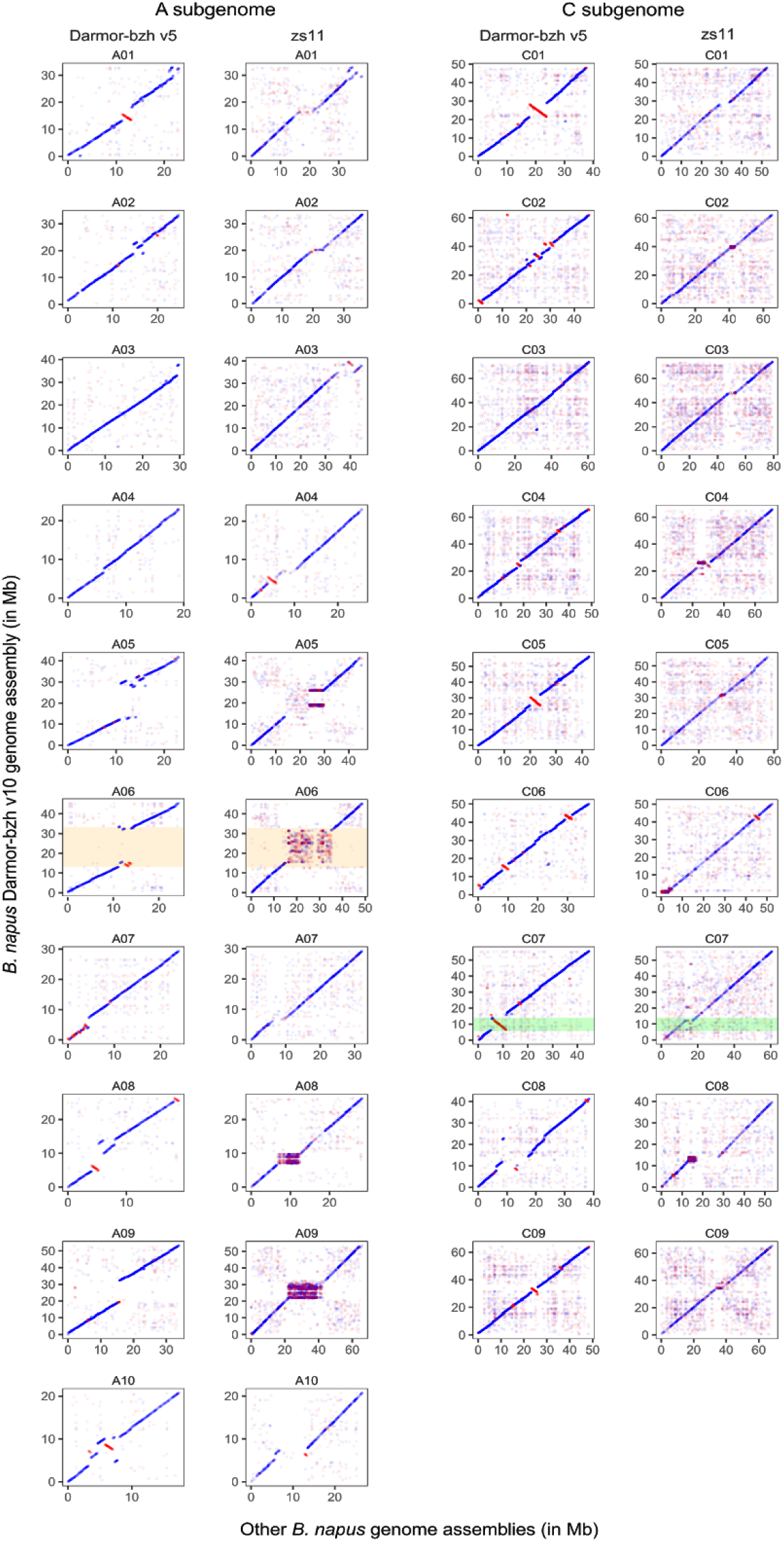
Genome-wide alignments of *B. napus* Darmor-bzh v10 (Y axis) with Darmor-bzh v5 and ZS11 genome assemblies (X axis). Each dot corresponds to syntenic regions of the genomes that are aligning with high confidence. Blue dots are corresponding to regions aligned in the correct orientation (+) while red dots represent regions aligned in an inverted orientation (-). The new assembly of a centromeric region in Darmor-bzh v10 compared to Darmor-bzh v5 is highlighted in orange for the A06 chromosome. An example of a large misassembled inverted region whose orientation has been corrected within the new Darmor-bzh v10 genome assembly is highlighted in green for the chromosome C07.

## Genome annotation

### Detection of modified bases

The nanopore technology has the advantage of reading native DNA molecules and can distinguish modified from natural bases because they affect the electrical current flowing through the pore differently [32]. We analysed the electric signal to detect the 5-methylcytosine (5mC) in the CpG context of the whole genome of *B. napus* Darmor-bzh. The nanopore reads were mapped using minimap2 [33] version 2.17-r941 with the ‘-x map-ont’ parameter. We called the 5mC and computed frequencies using nanopolish version 0.12.5 with a minimum log-likelihood threshold of 2. CpG positions showing a frequency above 50% and covered by at least 4 reads were considered methylated. The proportion of methylated CpG per 100Kb across the genome is shown in Figure 1.

### Correction of direct RNA reads

Direct RNA reads were corrected using TALC [34] with default parameters (except -k=21) and Illumina reads from leaves and roots samples of *B. napus* downloaded from the EBI database (ERX397788 and ERX397800). Before the correction step, raw reads were masked using DustMasker [35], and we kept reads with at least 150 unmasked positions and at least 75% of unmasked bases. From 10,416,515 input reads, we retained 9,099,437 nanopore reads and TALC successfully corrected 8,523,238 reads (93.7%) with an average length of 645bp (Table S9).

### Transposable elements annotation

Transposable elements (TE) were annotated using RepeatMasker [36] (with default parameters) and transposable element libraries from [8]. We masked near 54% of the genome and LTR Copia and Gypsy elements are the most abundant (15.5% and 13.0% respectively). The three Darmor-bzh assembly releases and the fourteen other *Brassica* genome assemblies were masked using the same procedure (Table 4).

### Gene prediction

Gene prediction was performed using several proteomes: eight from other genotype of *B. napus* [11] *(Westar, Zs11, QuintaA, Zheyou73, N02127, GanganF73, Tapidor3 and Shengli3), Arabidopsis thaliana* (UP000006548), the *B. napus* pan-annotation [37], resistance gene analogs (RGA) from *Brassica* [38] and the 2014 annotation of Darmor-bzh [8]. Low complexity in protein and genomic sequences were masked respectively with the SEG and DustMasker algorithms [35]. Proteomes were then aligned on the genome in a two-steps strategy. First, BLAT [39] (with default parameter) was used to fastly localize corresponding putative regions of these proteins on the genome. The best match and the matches with a score greater than or equal to 90% of the best match score have been retained. Second, alignments were refined using Genewise [40] (default parameters), which is more accurate for detecting intron/exon boundaries. Alignments were kept if more than 75% of the length of the protein is aligned on the genome.

In addition, we used the error-corrected direct RNA reads produced using the PromethION device. As already reported [41], the detection of splicing sites using raw reads is difficult. For example, on a subset of 1,000 reads, the proportion of GT-AG splicing sites is 6% lower in the raw reads compared to the corrected reads (Table S10). Error-corrected reads were aligned on the genome in a two-steps strategy. First, BLAT (with default parameters) was used to fastly localize corresponding putative regions of these RNA reads on the genome. The best match for each read was retained and a second alignment was performed using Est2Genome [42] (default parameters), to detect exons boundaries. From the 8,523,238 corrected reads, we retained 4,651,118 alignment that covered at least 80% of the read with at least 95% identity. To reduce the redundancy, we clustered the alignments on the genomic sequence using bedtools [31] and kept for each cluster the alignment with the highest score. The 70,904 resulting alignments (average identity of 97.5% and 4.07 exons per alignment) were used as input for the gene prediction.

All the transcriptomic and proteins alignments where combined using Gmove [43] which is a easy-to-use predictor with no need of pre-calibration step. Briefly, putative exons and introns, extracted from alignments, were used to build a graph, where nodes and edges represent respectively exons and introns. From this graph, Gmove extracts all paths from the graph and searches open reading frames (ORFs) which are consistent with the protein evidences. Finally, we decided to exclude single-exon genes predicted by RNA-Seq only and composed of more than 80% of untranslated regions (UTRs). Following this pipeline, we predicted 108,190 genes with 4.55 exons per gene in average.

Quality of the gene prediction was estimated using the single-copy orthologous gene analysis from BUSCO v4 [44] with Brassicales version 10 which contains 4,596 genes (Table 1).

### Localization and annotation of pericentromeric regions

We retrieved the markers flanking each centromere of *B. napu*s chromosomes [45] and aligned them against the different assembly versions of *B. napus* Darmor-bzh to determine the improvement of the highly repeated pericentromeric regions. The (peri)centromere localization was searched using six (peri)centromeric-specific repeat sequences of Brassica (CentBr1, CentBr2, CRB, PCRBr, TR238 and TR805) which were blasted (e-value: 10-6) against the Darmor-bzh v10 genome sequence using BLASTn [46] as described in [45]. The length and number of genes present in the different versions were compared.

### Annotation of the Resistance Genes Analogs

We identified the Resistance Genes Analogs (RGA) in Darmor-bzh v5 and v10 gene annotations using RGAugury [47] and the following command: ‘RGAugury.pl -p {peptides.fa} -e 1e-5 -c 10 -pfx {output}’. The latest versions of all the databases and tools used by RGAugury were downloaded as of April 2020 following the installation instructions provided in the RGAugury online repository [48].

Briefly, this pipeline enables to identify different major RGA families (NBS encoding; RLP: membrane associated receptor like proteins; RLK: surface localized receptor like protein kinases: RLK). Based on the presence or absence of some domain structures (CC: coiled-coil, TIR: Toll/Interleukin 1 Receptor, NB-ARC: nucleotide binding site activity regulated cytoskeleton; LRR: leucine rich repeat), the NBS encoding proteins were subdivided into eight different categories [49]. Finally all the putative RGA proteins identified in Darmor-bzh v5 were blasted against those identified in this new assembly in order to establish a correspondence between these two versions, but also to identify newly annotated RGA genes in Darmor-bzh v10.

## An improved version of the Darmor-bzh genome assembly

### Comparison with existing assemblies and annotations

The two published releases of Darmor-bzh [8,9] were generated using 454 and Illumina reads and have a low contiguity (Table 1). These fragmented assemblies were difficult to organize at the chromosome level, and as a result only 553Mb and 690Mb of sequences were respectively anchored on the 19 chromosomes. In comparison, the 19 chromosomes of the ONT assembly contains 849Mb. The gene completion (BUSCO score) of the first release and of this one are similar (97.7% and 98.6%), showing that long-read assemblies mainly impact the repetitive compartment of the genome. However 98.8% of the genes are now placed on their respective chromosomes rather than on unplaced scaffolds, in comparison only 80% of the predicted genes were located on pseudomolecules in Darmor-bzh v5 (Figure 3B). These improvements will greatly help the Brassica community to identify the genes underlying agronomic traits of interest found using quantitative genetics.

**Figure 3.**
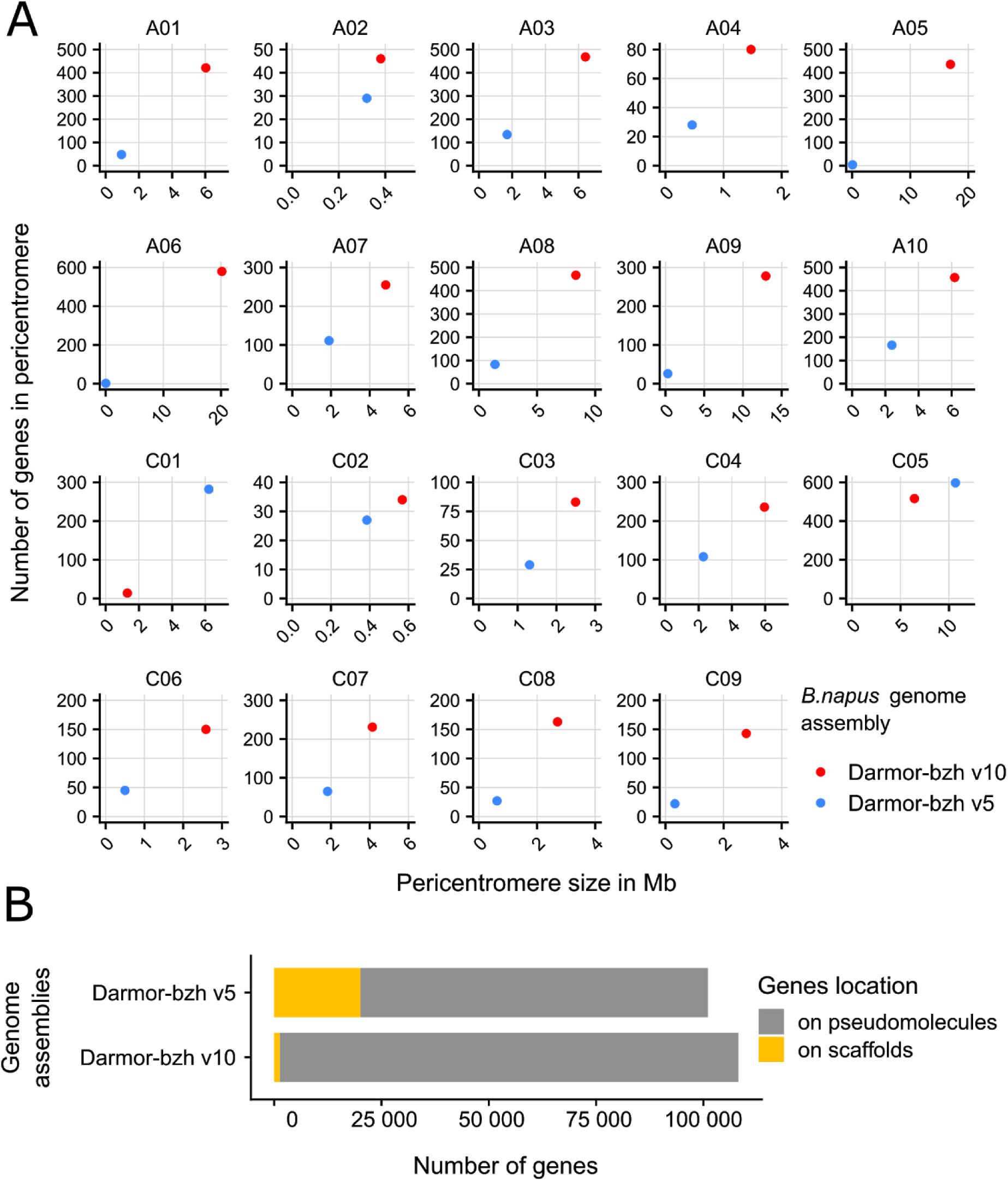
Improvements of the *B. napus* Darmor-bzh v10 genome assembly and annotation compared to Darmor-bzh v5 by the use of long reads sequencing. **A**. Genome-wide pericentromeres size (in Mb) and gene content using centromere flanking markers (Mason et al. 2016, refer to methods). The Darmor-bzh v10 genome assembly now harbours larger pericentromeres with more annotated genes than in the previous genome. **B**. Repartition of genes location on chromosome and unplaced scaffolds.

We compared the short- and long-read assemblies and observed large newly assembled regions comprising centromeric sequences (Figure 2, highlighted in orange on chromosome A06). By aligning the flanking markers of *B. napus* centromeres [45], we were able to identify the positions of the approximative pericentromeric regions in Darmor-bzh v5 and the present Darmor-bzh v10 assembly (Table S12 and Figure 1). Thereafter, we compared the lengths and gene contents of these regions between Darmor-bzh v5 and v10 (Figure 3A). On average, we observed an increase of sequence assembly in pericentromeric regions by 80 fold, with some extreme cases like the A06 pericentromere that is 1,180 times larger in Darmor-bzh v10 compared to Darmor-bzh v5. Concerning pericentromeric gene content, we show an average increase of 24.4 fold, with the A06 pericentromere showing 290 times more genes in Darmor-bzh v10 compared to Darmor-bzh v5. There was only two exceptions (C01 and C05), where we identified a decrease in pericentromere size and gene content in Darmor-bzh v10 compared to Darmor-bzh v5. However, these results may in fact be attributable to misassembled inverted regions in Darmor-bzh v5, which changed the position of the markers on which our detection of pericentromeric regions is based. Indeed, comparison of the different *B. napus* Darmor-bzh assemblies allowed to notably identify 27 large misassembled inverted regions (LMIs: superior to 700 Kb) in Darmor-bzh v5 that are now correctly assembled in the Darmor-bzh v10 assembly. These misassembled regions were validated using the new ZS11 assembly [11] as a control (Figure 2, highlighted in purple on the C07 chromosome). The 27 detected LMIs totalize 3,494 genes and 45Mb in Darmor-bzh v10, with the largest located on chromosome C01, measuring 6Mb and encompassing 276 genes. We provide a list of the large misassembled inverted regions detected in Darmor-bzh v5 and corrected in Darmor-bzh v10, alongside with the corresponding genes in these regions (Table S13).

All these analyses highlight the improvements of the Darmor-bzh v10 genome assembly thanks to the use of ONT long-reads sequencing. Overall, this high-quality genome assembly of *B. napus* Darmor-bzh will be particularly useful to the *Brassica* community to decipher genes underlying agricultural traits of interest.

### Comparison of available Brassica genome assemblies

We downloaded the 14 available *Brassica* long-read assemblies (and annotations) and computed usual metrics. For each assembly, contigs were generated by splitting sequences at each gap and the gene completion was evaluated on predicted genes using BUSCO [44] and the conserved genes from the brassicales database (N=4,596 genes). Not surprisingly the gene content is high, between 97.9% and 98.8% for all the ten *B. napus* assemblies (Table 2), and is lower for the diploid species (due to the presence of a single-genome), except for *B. rapa* Chiifu that has a surprising number of genes (>60k). Likewise, the repetitive content of these ten genomes is similar, between 54% for Darmor-bzh and 60% for no2127. Concerning the diploid genomes and as already reported [50], *B. oleracea* (C) genome assemblies contain more repetitive elements than the *B. rapa* (A) and *B. nigra* (B) genome assemblies (Table 4).

The main observed differences are the contig length that affects the number of gaps in each assembly (from 268 gaps for Darmor-bzh to 5,460 gaps for Zs11) and the proportion of anchored sequence (from 765 Mb for Express617 to 961 Mb for Zs11). We further investigated these two differences that we believe to be related to the technologies used.

We observed a significant difference in contiguity when comparing ONT and PACBIO assemblies. The eleven *Brassica* genomes sequenced using PACBIO have a contig N50 between 1.4Mb and 3.6Mb whereas all the ONT assemblies have a contig N50 higher than 5.5Mb. The ability of the nanopore technology to sequence large fragments of DNA appears to be an advantage for assembling complex genomes. In the case of Darmor-bzh, we were able to obtain more than 50,000 reads longer than 100Kb (representing 6X of coverage).

This dataset allowed us to generate a contiguous assembly with only 268 gaps. As a comparison, we found 5,460 in the PACBIO assembly of the Zs11 genotype. This difference is observed in all the A and C genomes and subgenomes that have been sequenced using long reads, and may be directly related to the longer length of ONT reads (Figure 4). Although the number of gaps is higher in the PACBIO assemblies, the total number of N’s is lower at least for the assemblies that have been organized with Hi-C data. Indeed, the Hi-C pipelines generally order contigs and add a fixed gap size between two contigs. We investigated the 500bp gaps in the Zs11 assembly and aligned their flanking regions on the Darmor-bzh assembly using blat [39]. We applied stringent criteria (alignment on the same chromosome, score > 4000, alignment of at least 1 Kb of each of the flanking regions and alignment covering less than 100Kb on the Darmor-bzh assembly) and we found a location for 367 of the 5,460 gaps of Zs11. These 367 regions covered more than 19Mb of the Darmor-bzh assembly with an average size of 52 Kb. Even if these regions could be different between the two genotypes, we examined their content in transposable elements and found a high proportion of bases annotated as repetitive elements (82.9%) and a different distribution of the classes of elements compared to the whole genome (Table S11). We observed a higher proportion of Copia elements, but especially LINE and Satellite elements. For example, we observed large LINE elements (>50Kb) as shown in Figure 5, where ultralong nanopore reads covered this element and avoided a contig breakage in this area.

**Figure 4.**
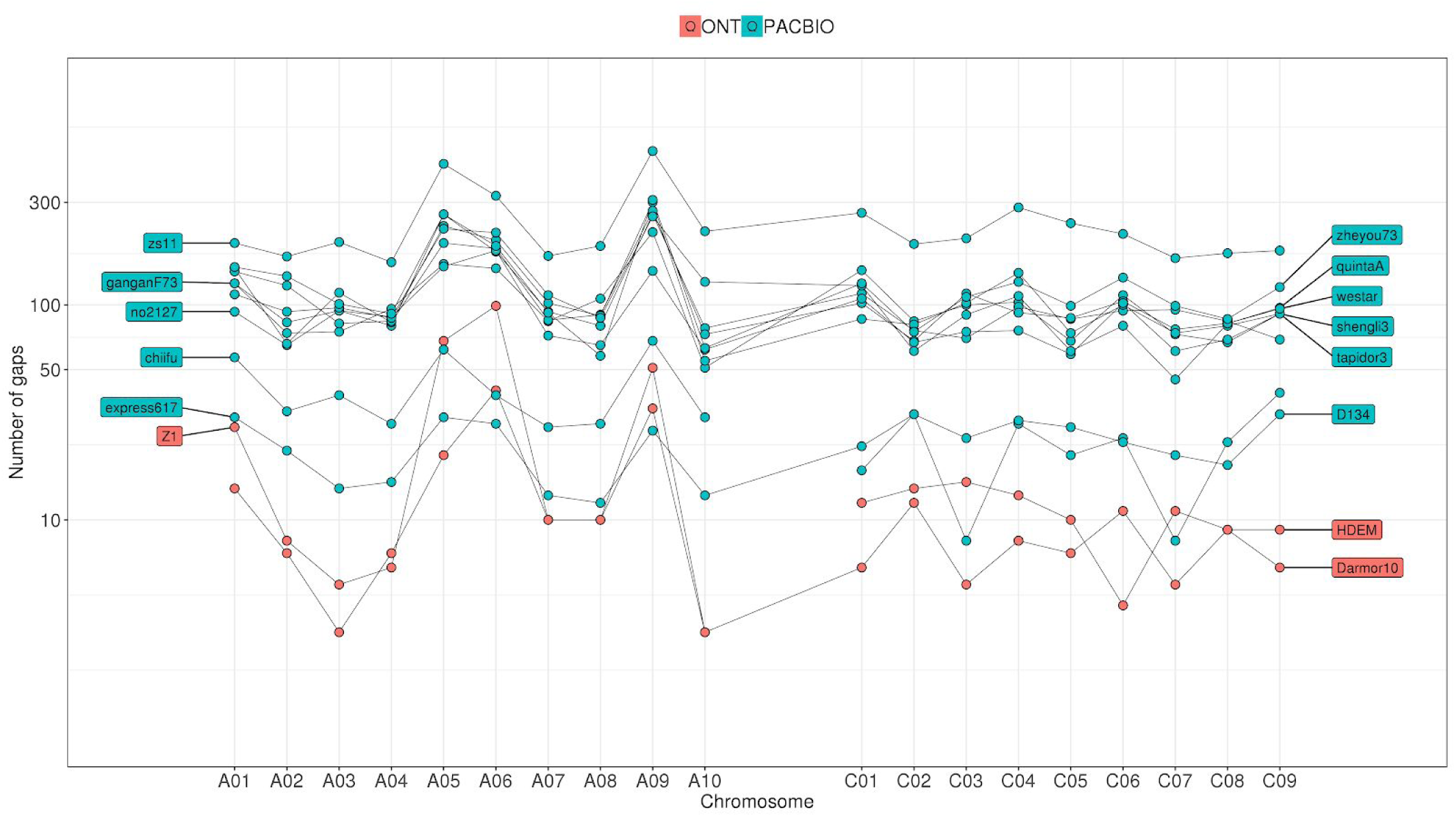
Distribution of the number of gaps per chromosome in *Brassica* genomes. Each dot represents the number of gaps in a given chromosome and genome assembly. PACBIO assemblies are in blue and ONT assemblies in orange.

**Figure 5.**
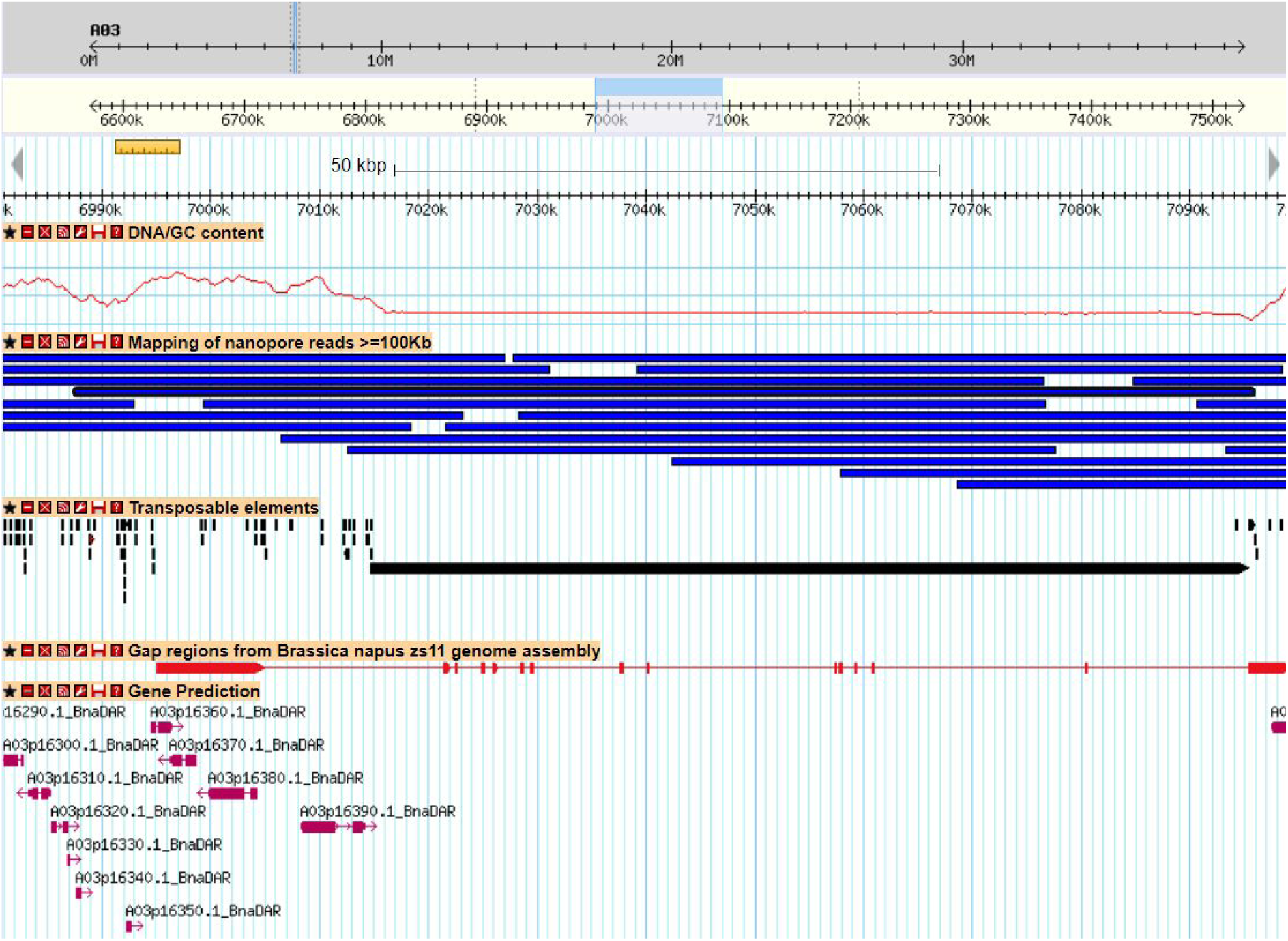
Example of a genomic region of *B. napus* Darmor-bzh assembly that corresponds to a region that contain a gap in the Zs11 genome assembly. First track represents the GC content. ONT reads (longer than 100Kb) are in the second track (blue boxes), and a 115 Kb read that spanned the whole region is surrounded by a black box. Transposable elements are shown as black boxes, and the region absent from Zs11 contain a large LINE element (80,707 bp). The alignment of the Zs11 sequence (20Kb around the 500bp gap) is represented by red boxes, with thin red lines representing missing sequences in the Zs11 genome assembly that are present in Darmor-bzh v10 (gaps). Predicted genes are in the last track, in purple.

Interestingly, A chromosomes seem more difficult to assemble (Figure 4). Even if A genome is smaller than C genome and contains less transposable elements (Table 4), the number of contigs (and consequently the number of gaps) in A genome, or subgenome, is twice the number of contigs in C genome, or subgenome (gap track on Figure 1 and Figure 4). Chromosomes A05, A06 and A09 appear to be the most challenging to assemble due to the high content of transposable elements in their centromeric regions. They alone contain 43% of the gaps (and 51% of the undetermined bases) present on the 19 chromosomes. However these chromosomes have been shown to be highly variable in length across *B. rapa* genotypes [51].

The second difference observed between all these assemblies is the proportion of anchored sequence which is higher for the genome assembly that has been organized using Hi-C data (95.0%, Zs11) although it is the most fragmented (Table S8). Indeed the organization of small contigs is complicated using optical or genetic maps because it requires a sufficient number of restriction sites or markers. This aspect probably explains the small proportion of the Express617 assembly (82.7%) that has been anchored on the 19 chromosomes. The other genomes have been organized using comparative genomics (synteny with the Zs11 assembly). Although it is a convenient method, it may generate false organization and the proportion of anchored bases is variable, from 89.1% to 92.7%, depending on the conservation between the two genomes. We think that if the assembly is highly contiguous, optical maps have the important advantage of estimating the size of gaps, which remains a limitation when using Hi-C data.

## Sequencing of native RNA molecules

The sequencing of RNA is traditionally performed using the Illumina technology where the RNA molecules are isolated and then reverse transcribed (RT) to cDNA that is more stable and allow the amplification prior to the sequencing. The Oxford Nanopore technology is the first to propose the sequencing of native RNA molecules without the RT step. One of the main advantage is a better quantification of gene expression when compared to methods that requires a RT [41]. Moreover the sequencing of native RNA molecules preserves modified bases and theoretically allows to detect them [52,53]. Here, we sequenced a mix of leaf and root samples using the PromethION device and generated 10,416,515 reads with an average size of 559 bp (Table S9). This dataset was corrected using TALC as described previously and corrected reads were used in the gene prediction workflow.

### Alternative splicing events

Independently, we detected alternative splicing (AS) events, as skipping exon and intron retentions. Aligning noisy reads makes it difficult to accurately detect splice sites and therefore 5’ and 3’ alternative splice sites. From the genomic alignments of raw reads, with an identity percent higher than 90%, we used bedtools to extract skipping exon (reads with one exon that is entirely covered by an intron, smaller than 3Kb to avoid mapping errors) and intron retentions (reads with one exon that fully spanned at least two consecutive exons). We detected intron retentions and skipping exon in respectively 16% and 3% of the 108,190 annotated genes. For example, by inspecting mutually exclusive exons, we were able to find an already described event that is conserved between *Brassica* and *Arabidopsis thaliana* [54] (Figure 6). It is obvious that the sequencing of RNA using long-reads will allow to detect co-occurrence of splicing events that is difficult if not impossible to do with short-reads.

**Figure 6.**
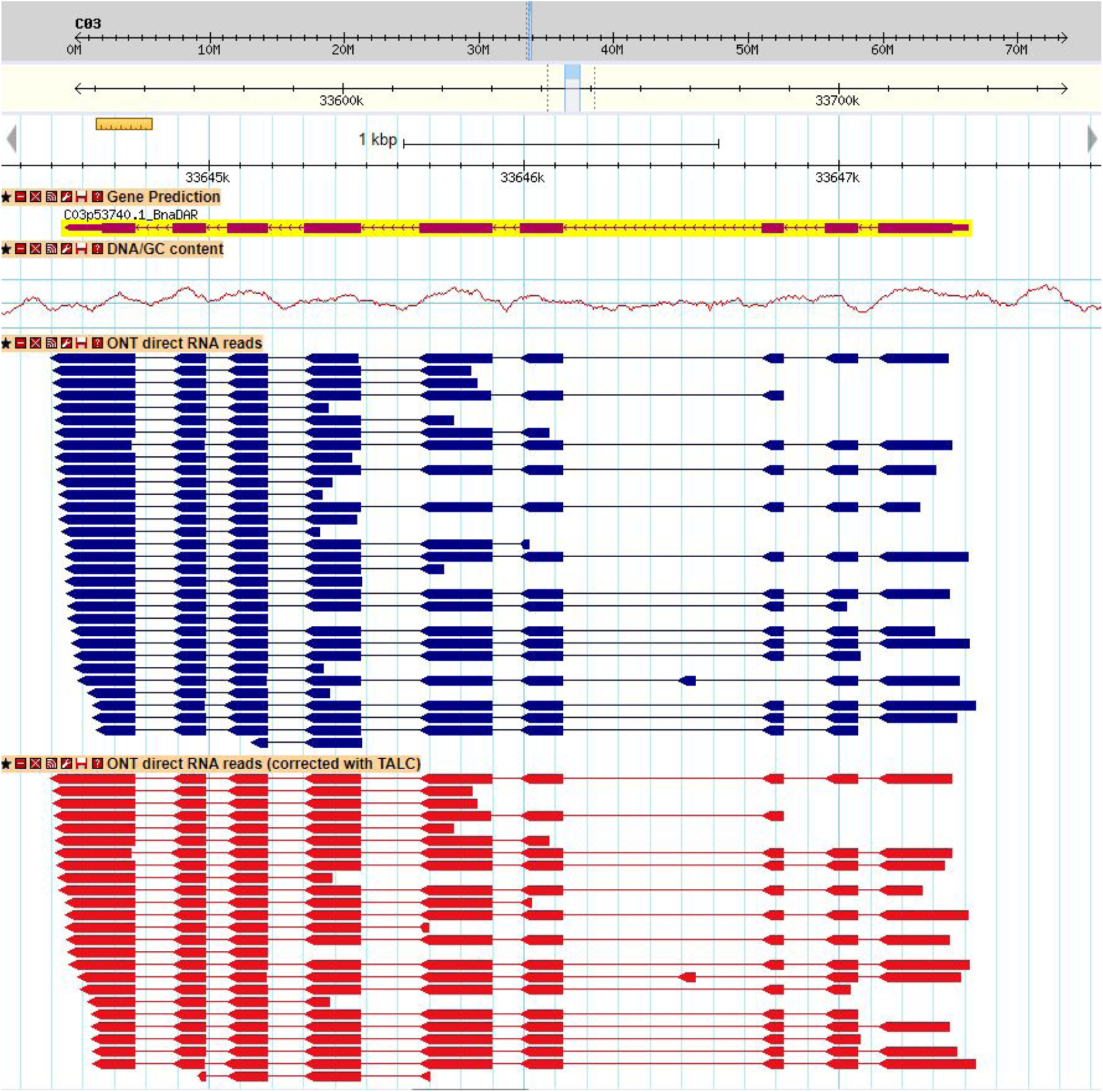
Example of a splicing event detected using long reads. The exon in the third introns of the gene prediction (C03p53740.1_Bna_DAR, Gene prediction track) is mutually exclusive with the third exon. This alternative splice form is detected by a single nanopore read (blue track), and maintained by the TALC error-correction (red track).

### Comparison of the Resistance Genes Analogs catalogs

Using the RGAugury pipeline [47], we annotated 2,788 and 2,952 Resistance Genes Analogs (RGA) in the gene catalogs of Darmor-bzh v5 and v10 respectively (Tables S14 and S15). In Darmor-bzh v5 only 83.5% of the RGAs were located on anchored scaffolds, while we identify 99.4% of the RGAs on chromosomes in Darmor-bzh v10. In average 153±46 RGAs are located on each pseudomolecule in Darmor-bzh v10 compared to 122±33 in Darmor-bzh v5 (rounded mean±standard deviation). Overall, more RGAs were annotated in the Darmor-bzh v10 annotations in almost all categories, thanks to the long-reads sequencing. Most of the RGAs found in Darmor-bzh v5 were conserved in Darmor-bzh v10 (81,4%, Table 3 and Figure 7A). However, we observed that 108 RGAs annotation shifted from one RGA class to another between the two versions due to the identification of additional domains in these proteins (Figure 7A). Between the Darmor-bzh v5 and Darmor-bzh v10 annotations, the CN and NL RGA classes shifted the most to CNL (CC, NB-ARC and LRR domains, Figure 7B and 7C), which are displaying one additional domain compared to CN (CC and NB-ARC domains) or NL (NB-ARC and LRR domains) RGAs. In addition, we identified dozens of examples were few adjacent RGA genes in the Darmor-bzh v5 now corresponds to a single gene in the Darmor-bzh v10.

**Table 3.** Resistance Genes Analogs categories and repartition between chromosomes and unplaced scaffolds in *B. napus* Darmor-bzh v5 and v10 genome assemblies.

| Annotation | RGA Location | RGA classes |  |  |  |  |  |  |  |  |  |  | Total |
| --- | --- | --- | --- | --- | --- | --- | --- | --- | --- | --- | --- | --- | --- |
|  |  | NBS | CNL | TNL | CN | TN | NL | TX | Others | RLP | RLK | TM-CC |  |
| Darmor-bzh<br>v5 RGA | genome-wide | 37 | 53 | 164 | 33 | 60 | 80 | 143 | 24 | 274 | 1502 | 418 | 2788 |
|  | pseudomolecules | 27 | 45 | 144 | 27 | 54 | 67 | 112 | 22 | 231 | 1259 | 340 | 2328 |
|  | scaffolds | 10 | 8 | 20 | 6 | 6 | 13 | 31 | 2 | 43 | 243 | 78 | 460 |
| Darmor-bzh<br>v10 RGA | genome-wide | 45 | 71 | 189 | 22 | 58 | 85 | 128 | 29 | 285 | 1569 | 471 | 2952 |
|  | pseudomolecules | 45 | 71 | 184 | 22 | 57 | 84 | 128 | 29 | 282 | 1561 | 471 | 2934 |
|  | scaffolds | 0 | 0 | 5 | 0 | 1 | 1 | 0 | 0 | 3 | 8 | 0 | 18 |

**Table 4.** Repetitive element content of the Brassica long-reads assemblies (ten polyploid *B. napus*, two diploid *B. rapa*, one diploid *B. nigra* and two diploid *B. oleracea* genomes).

|  | Total length (Mb) | Masked proportion | LTR Copia | LTR Gypsy | DNA CMC-EnSpm | LINE | RC Helitron | DNA TcMar | DNA MULE-MuDR |
| --- | --- | --- | --- | --- | --- | --- | --- | --- | --- |
| Darmor-bzh | 924 | 53.86% | 15.55% | 13.05% | 6.66% | 6.37% | 5.26% | 2.05% | 1.93% |
| Westar | 1,008 | 58.91% | 16.88% | 13.38% | 6.26% | 6.92% | 4.87% | 1.92% | 1.77% |
| Express617 | 925 | 56.32% | 15.92% | 12.46% | 6.54% | 6.63% | 5.22% | 2.05% | 1.88% |
| Tapidor3 | 1,014 | 59.01% | 16.49% | 13.53% | 6.17% | 7.05% | 4.86% | 1.91% | 1.76% |
| Shengli3 | 1,002 | 58.50% | 16.15% | 13.16% | 6.16% | 6.97% | 4.86% | 1.92% | 1.78% |
| QuintaA | 1,004 | 58.60% | 16.40% | 13.23% | 6.29% | 7.03% | 4.93% | 1.93% | 1.80% |
| No2127 | 1,012 | 59.85% | 17.08% | 13.74% | 6.23% | 7.02% | 4.77% | 1.91% | 1.72% |
| GanganF73 | 1,034 | 58.76% | 16.60% | 13.48% | 6.23% | 6.86% | 4.82% | 1.91% | 1.75% |
| Zheyu73 | 1,016 | 59.06% | 16.49% | 13.59% | 6.28% | 7.04% | 4.87% | 1.91% | 1.79% |
| Zs11 | 1,010 | 58.62% | 16.59% | 13.37% | 6.36% | 6.92% | 4.88% | 1.95% | 1.78% |
| Z1 | 402 | 46.95% | 13.22% | 13.23% | 2.88% | 5.16% | 4.37% | 1.39% | 1.53% |
| Chiifu | 353 | 48.46% | 12.89% | 13.18% | 3.29% | 5.60% | 5.15% | 1.63% | 1.82% |
| NI100 | 506 | 53.70% | 16.51% | 22.59% | 4.08% | 4.93% | 3.17% | 1.42% | 1.79% |
| D134 | 530 | 57.88% | 15.61% | 13.23% | 9.16% | 6.24% | 5.62% | 2.56% | 2.08% |
| HDEM | 555 | 57.14% | 16.06% | 12.68% | 8.74% | 6.31% | 5.42% | 2.46% | 1.99% |

**Figure 7.**
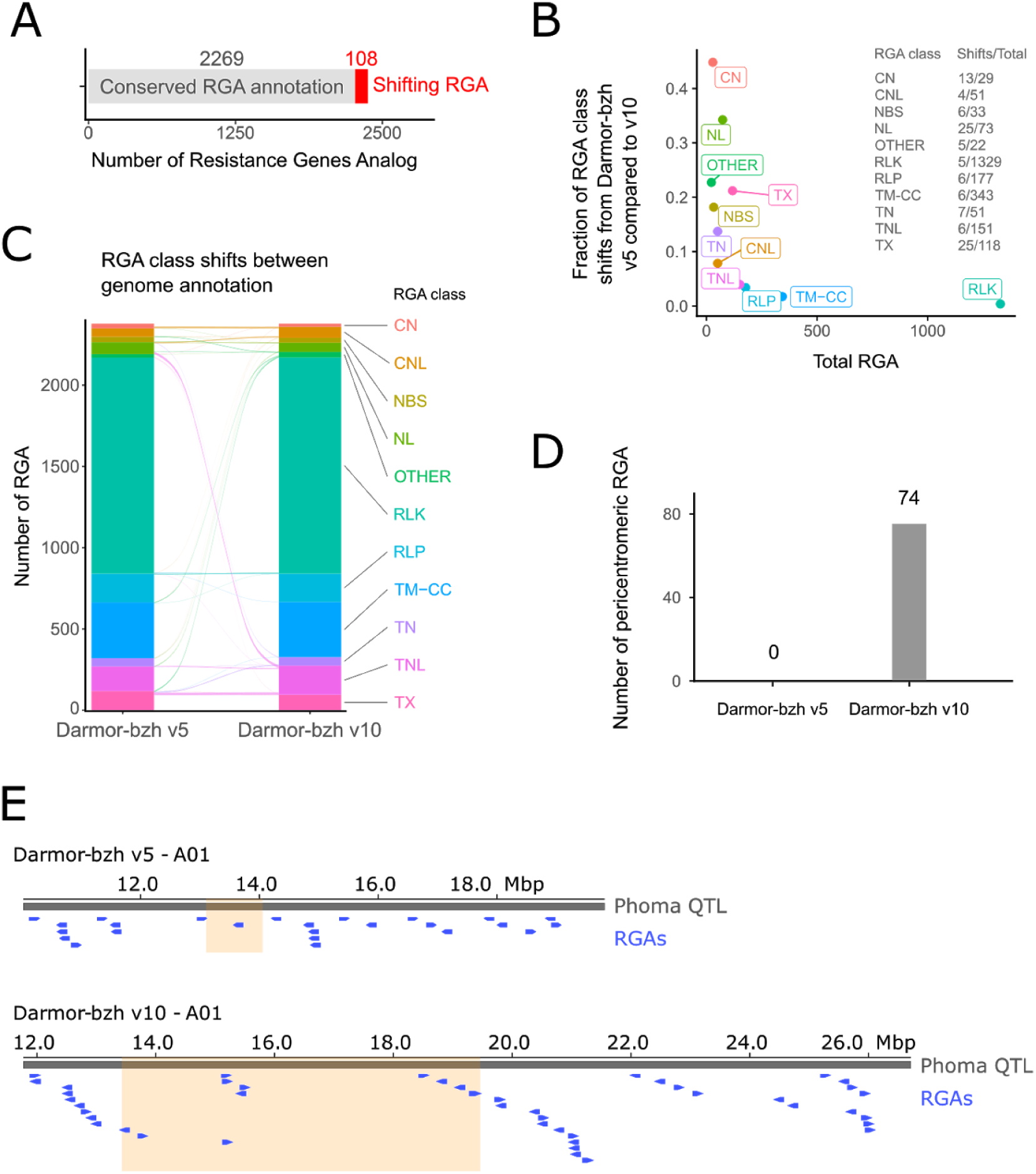
Resistance Genes Analogs (RGA) annotation improvement in *B. napus* Darmor-bzh v10 using long reads direct RNA sequencing. **A**. Conservation of the RGA categories between Darmor-bzh v5 and v10 genome annotations. Shifting RGAs are genes whose sequence is matching, but that have been annotated in a different RGA class by RGAugury between the v5 and v10 genome annotations. Such shifting RGAs only represent a small fraction of the annotated RGAs. **B**. Total number of RGA plotted against the ratio of shifting RGAs over total RGAs by RGA class. CN and NL RGAs are the categories whose annotation shifted the most between Darmor-bzh v5 and v10. A detail of the raw data used to compute the ratio is available on the right part of the plot. **C**. Detail of the RGA class shifts between Darmor-bzh v5 and v10 made using ggalluvial [55]. The size of the arcs is proportional to the number of shifting RGAs from one RGA class to another between the two annotations. **D**. Number of pericentromeric RGAs found in Darmor-bzh v10 compared to the v5 genome annotation. **E**. Genome browser snapshot using pyGenomeTracks [56] of the phoma canker resistance QTL (grey) in Darmor-bzh v5 (top) and Darmor-bzh v10 (bottom). RGA genes (blue) and pericentromeric regions (orange) are displayed.

As pericentromeric region assembly was greatly improved in Darmor-bzh v10, we compared the number of RGA in such regions between the two assemblies and found 74 RGAs in Darmor-bzh v10 compared to no RGAs in Darmor-bzh v5 (Figure 7D). To further highlight the interest of assembling such regions using ONT technologies, we compared the size and number (including RGA) of a resistance QTL to phoma stem canker overlapping the A01 centromere [47,50]. In this new assembly, we were able to identify 45 candidate RGA compared to 29 in Darmor-bzh v5 (Figure 7E). This finding opens new ways to understand the mechanism underlying this QTL.

## Conclusions

In this study, we generated the most contiguous *B. napus* genome assembly thanks to the ONT long reads and Bionano optical maps. In addition, the ONT dataset allow to detect modified bases, such as C5-methylcytosine (5mC), without any specific library preparation. We have shown that the generation of ultra-long read is a game-changer for assembling complex regions, composed of transposable elements, common in plant genomes. Consequently, the combination of ultralong reads and optical maps is today a method of choice for generating assembly of complex genomes. Without forgetting that the nanopore technology is in constant evolution, in particular base calling softwares, which make it possible to improve an assembly starting from existing data. In addition, we have predicted genes in this new assembly using direct RNA sequencing and have used this original dataset to detect splicing events and show that this technology can be used to discover complex events, such as the co-occurence of events as mutually exclusive exons. This improved version of the *B. napus* Darmor-bzh reference genome and annotation will be valuable for the Brassica community, which has now been working with the short-read version for almost six years, particularly to decipher genes underlying agricultural traits of interest.

## Additional files

All the supporting data are included in five additional files which contain a) Tables S1-S11 and Figure S1, b) Table S12, c) Table S13, d) Table S14 and e) Table S15.

Acknowledgements

This work was supported by the Genoscope, the Commissariat à l’Energie Atomique et aux Energies Alternatives (CEA) and France Génomique (ANR-10-INBS-09-08). We acknowledge the BrACySol BRC (INRA Ploudaniel, France) that provided us with the seeds of *B. napus* cv. Darmor-bzh. The authors are grateful to Oxford Nanopore Technologies Ltd for providing early access to the PromethION device through the PEAP, and we thank the staff of Oxford Nanopore Technology Ltd for technical help.

## Availability of supporting data

Darmor-bzh is a French winter variety, freely available in Plant Genetic Resources BrACySol (Rennes, France). The Illumina and PromethION sequencing data and the Bionano optical maps are available in the European Nucleotide Archive under the following projects PRJEB39416 and PRJEB39508. The genome assembly, gene predictions and a genome browser are freely available at http://www.genoscope.cns.fr/plants.

## Author’s contributions

CF, GD and CC extracted the DNA and the RNA. CC and AL optimized and performed the sequencing and the optical maps. CF, RD and FB generated the different genetic maps for Brassica napus. BI, CB and JMA performed the genome assembly. CF, JM, JFdC, FB, MRG, CB and JMA performed the anchoring of the *B. napus* scaffolds. SE performed the detection of modified bases. CDS performed the error correction of direct RNA reads and the detection of alternative splicing events. JMA and CDS performed the gene prediction of the genome assembly. GR, MRG, JB, LM, BI, CB, CDS and JMA performed the bioinformatic analyses. MRG, GR, BI, CB and JMA wrote the article. MRG, AMC, FD, PW and JMA supervised the study.

