## Supplementary Data for "Long-reads assembly of the *Brassica napus* reference genome, Darmor-bzh"

**Supplementary Table 1.** Statistics of the sequencing dataset for *B. napus* Darmor-bzh

| | ONT (6 PromethION runs) | | | Illumina |
|---|---|---|---|---|
| | Raw reads | Longest reads | Filtlong reads | 1.5 lane of HS2500 Rapid Run |
| Cumulative size | 111.9 Gb | 36.1 Gb | 36 Gb | 97 Gb |
| # of reads | 15,831,026 | 462,512 | 712,158 | 215,472,682 |
| Coverage (estimated genome size 1.2Gb) | 93X | 30X | 30x | 81X |
| Coverage Reads >100Kb (estimated genome size 1.2Gb) | 5.75X | 5.75X | 1.97X | NA |
| N50 (bp) | 42,806 | 76,025 | 55,397 | NA |
| Longest (bp) | 1,084,288 | 1,084,288 | 994,661 | 250 |
| Accession numbers | | NA | NA | |

**Supplementary Table 2.** *B. napus* Darmor-bzh ONT assembly statistics.

| Assembler | Subset of reads used | Cumulative Size | # contigs | N50 (L50) | N90 (L90) | CPU time (h) |
|---|---|---|---|---|---|---|
| Smartdenovo | All reads | 916,568,779 | 731 | **10,218,000** (31) | 597,059 (156) | 7,483 |
| | Filtlong reads | 835,563,225 | **625** | 4,161,078 (53) | 597,575 (238) | 2,079 |
| | Longest reads | 819,736,553 | 773 | 2,041,018 (107) | 448,924 (428) | 2,863 |
| Redbean | All reads | 1,870,584,980 | 53,028 | 63,466 (8,388) | 14,710 (31,095) | 243 |
| | Filtlong reads | 1,476,953,275 | 27,745 | 109,469 (3,407) | 23,825 (14,495) | **176** |
| | Longest reads | 1,273,533,804 | 21,319 | 127,500 (2,252) | 127,500 (2,252) | 184 |
| Flye | All reads | - | - | - | - | crashed |
| | Filtlong reads | 927,471,373 | 1,516 | 8,067,213 (34) | **616,556** (165) | 1,632 |
| | Longest reads | **937,929,502** | 1,594 | 10,072,792 **(29)** | 561,876 **(146)** | 1,472 |

**Supplementary Table 3.** *B. napus* Darmor-bzh bionano dataset (raw molecules)

| | BspQI molecules | DLE-1 molecules |
|---|---|---|
| Number of molecules | 2,124,074 | 16,646,860 |
| Total length (Mb) | 223,164 | 1,340,608 |
| Average length (Kb) | 105 | 81 |
| Molecule N50 (Kb) | 178 | 101 |
| Label density (/100Kb) | 7.7 | 16.6 |
| Number of flowcells | 2 | 1 |

**Supplementary Table 4.** *B. napus* Darmor-bzh bionano dataset (filtered molecules)

| | BspQI molecules | DLE-1 molecules |
|---|---|---|
| Number of molecules | 402,272 | 1,699,106 |
| Total length (Mb) | 101,656 | 397,015 |
| Average length (Kb) | 253 | 234 |
| Molecule N50 (Kb) | 257 | 229 |
| Label density (/100Kb) | 9.4 | 16.9 |

**Supplementary Table 5.** *B. napus* Darmor-bzh optical maps

| | BspQI map | DLE-1 map |
|---|---|---|
| Number of maps | 868 | 178 |
| Total map length (Mb) | 1,042 | 968 |
| Map N50 (Mb) | 1.7 | 18.2 |

**Supplementary Table 6.** *B. napus* Darmor-bzh hybrid scaffolding and polishing

| | ONT polished assembly (contigs) | Optical maps integration (scaffolds) | BiSCoT (contigs) | BiSCoT (scaffolds) |
|---|---|---|---|---|
| Cumulative size (bp) | 929,388,171 | 957,271,942 | 907,852,133 | 954,769,412 |
| # sequences | 1,869 | 1,432 | 1,581 | 1,378 |
| N50 (L50) | 9,972,764 (29) | 22,450,564 (15) | 11,241,391 (25) | 22,666,588 (15) |
| N90 (L90) | 553,072 (146) | 2,973,865 (53) | 718,812 (111) | 3,083,366 (52) |
| Maximum size (bp) | 34,824,884 | 48,039,605 | 46,624,681 | 47,917,037 |
| Number of N's (%) | 0% | 28,143,795 (2.94 %) | 0% | 25,966,328 (2.72%) |

**Supplementary Table 7.** *B. napus* Darmor-bzh chromosomal organization

| | 19 chromosomes | Unanchored scaffolds | A genome | C genome |
|---|---|---|---|---|
| Cumulative size (bp) | 866,915,903 | 56,879,860 | 346,465,994 | 520,449,909 |
| # sequences | 19 | 218 | 10 | 9 |
| N50 (L50) | 53,549,824 (7) | 652,631 (15) | 39,685,748 (4) | 62,297,340 (4) |
| # contigs | 505 | - | 157 | 78 |
| Contig N50 (L50) | 11,486,274 (24) | - | 9,804,805 (11) | 15,347,254 (12) |
| N90 (L90) | 29,390,524 (16) | 93,503 (142) | 23,101,716 (9) | 48,239,360 (7) |
| Maximum size (bp) | 73,669,886 | 4,834,143 | 53,549,826 | 73,669,886 |
| Number of N's (%) | 17,515,951 (2.20%) | 8,475,880 (14.90%) | 12,445,409 (3.59%) | 5,070,542 (0.97%) |

**Supplementary Table 8.** Statistics of the *B. napus* long-reads assemblies (ordered by contigs N50 value)

| | **genotype** | Long-read technology | Contig N50 (L50) | # gaps | Average gaps size (bp) | Long-range technology | Number of anchored bases | % of anchored bases |
|---|---|---|---|---|---|---|---|---|
| *B. napus* | darmor-bzh | ONT | 11,486,274 (24) | 268 | 96,984 | Optical maps | 867 Mb | 93.8% |
| *B. napus* | westar | PACBIO | 3,130,520 (93) | 2,165 | 500 | Comp | 935 Mb | 92.7% |
| *B. napus* | express617 | PACBIO | 3,002,211 (78) | 661 | 1,195 | Optical maps | 765 Mb | 82.7% |
| *B. napus* | tapidor3 | PACBIO | 2,855,025 (104)) | 1,771 | 500 | Comp | 921 Mb | 90.8% |
| *B. napus* | shengli3 | PACBIO | 2,825,656 (104) | 2,241 | 500 | Comp | 908 Mb | 90.6% |
| *B. napus* | quintaA | PACBIO | 2,801,289 (104) | 2,132 | 500 | Comp | 920 Mb | 91.6% |
| *B. napus* | no2127 | PACBIO | 2,704,645 (98) | 2,074 | 500 | Comp | 910 Mb | 89.9% |
| *B. napus* | ganganF73 | PACBIO | 2,696,026 (98) | 1,976 | 500 | Comp | 922 Mb | 89.1% |
| *B. napus* | zheyou73 | PACBIO | 2,103,870 (126) | 2,428 | 500 | Comp | 907 Mb | 89.2% |
| *B. napus* | zs11 | PACBIO | 1,506,624 (186) | 5,460 | 500 | Hi-C | 961 Mb | 95.0% |

**Supplementary Table 9.** Statistics of direct RNA nanopore reads.

| | **Raw reads** | Filtered reads | Error corrected reads (TALC) |
|---|---|---|---|
| Number of reads | 10,416,515 | 9,099,437 | 8,523,238 |
| Cumulative size (bp) | 5,819,930,143 | 5,673,021,329 | 5,497,587,252 |
| Average size (bp) | 559 | 623 | 645 |
| N50 (bp) | 737 | 745 | 763 |
| % of reads (>1Kb) | 12.3% | 13.9% | 14.9% |

**Supplementary Table 10.** Mapping of a sample of 1,000 reads using blat and est2genome.

| | Filtered reads | Error corrected reads (TALC) |
|---|---|---|
| % of mapped reads | 99.45% | 99.78% |
| Identity percent | 90.8% | 97.80% |
| Number of exons per model | 2.04 | 2.05 |
| % of GT-AG splice sites | 90.43% | 96.51% |

**Supplementary Table 11.** Repetitive content of the Darmor-bzh genome assembly compared to the regions present in Darmor-bzh but absent from zs11.

| | Total length (Mb) | Masked proportion | LTR Copia | LTR Gypsy | DNA CMC-EnSpm | LINE | Satellite |
|---|---|---|---|---|---|---|---|
| darmor-bzh | 924 | 53.86% | 15.55% | 13.05% | 6.66% | 6.37% | 1.66% |
| zs11 gaps | 19.2 | **82.89%** | **25.29%** | 13.69% | 5.15% | **16.64%** | **19.22%** |

**Supplementary Figure 1.** Example of a transcript isoform difficult to detect using short-reads (orange track). Two nanopore long-reads (blue track) show a co-occurence of the presence of the second exon with the absence of the fourth exon.

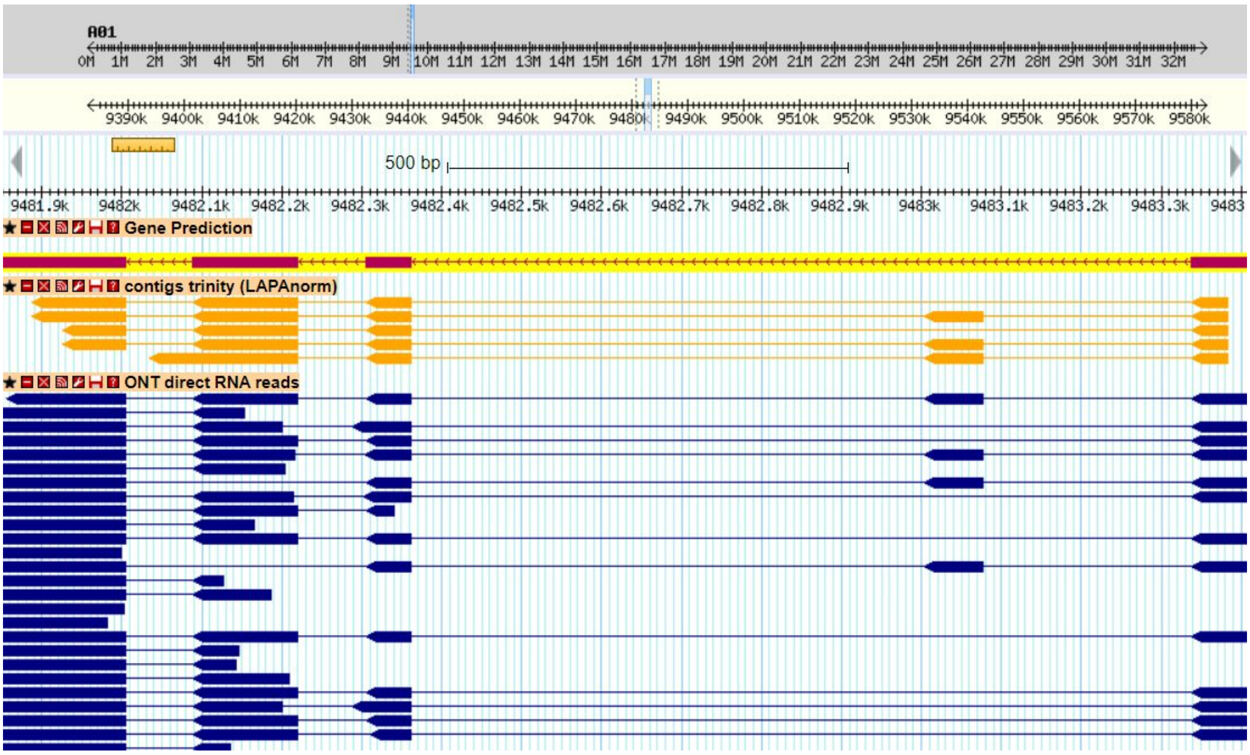
